# CUT&RUN: Targeted *in situ* genome-wide profiling with high efficiency for low cell numbers

**DOI:** 10.1101/193219

**Authors:** Peter J. Skene, Steven Henikoff

**Affiliations:** Howard Hughes Medical Institute, Basic Sciences Division, Fred Hutchinson Cancer Research Center, 1100 Fairview Ave N, Seattle, Washington, USA 98109

**Keywords:** transcription factors, histone modifications, epigenomic profiling, DNA sequencing

## Abstract

Cleavage Under Targets and Release Using Nuclease (CUT&RUN) is an epigenomic profiling strategy in which antibody-targeted controlled cleavage by micrococcal nuclease releases specific protein-DNA complexes into the supernatant for paired-end DNA sequencing. As only the targeted fragments enter into solution, and the vast majority of DNA is left behind, CUT&RUN has exceptionally low background levels. CUT&RUN outperforms the most widely-used Chromatin Immunoprecipitation (ChIP) protocols in resolution, signal-to-noise, and depth of sequencing required. In contrast to ChIP, CUT&RUN is free of solubility and DNA accessibility artifacts and can be used to profile insoluble chromatin and to detect long-range 3D contacts without cross-linking. Here we present an improved CUT&RUN protocol that does not require isolation of nuclei and provides high-quality data starting with only 100 cells for a histone modification and 1000 cells for a transcription factor. From cells to purified DNA CUT&RUN requires less than a day at the lab bench.

## INTRODUCTION

### Development of the protocol

All of the cells in a multicellular organism have the same genomic sequence, but different gene expression patterns underpin tissue specification. Differences in gene expression arise from the binding of transcription factors (TFs) and their recruitment of chromatin-associated complexes that modify and mobilize nucleosomes. As a result, genome-wide mapping of TFs, chromatin-associated complexes and chromatin states, including histone variants and post-translational modifications (PTMs), has become a major focus of research. For over 30 years, chromatin immunoprecipitation (ChIP) has been the predominant method of mapping protein-DNA interactions. With ChIP, cells are crosslinked with formaldehyde, then the entire cellular content is solubilized to fragment the chromatin fiber, and an antibody is added to isolate the chromatin fragments of interest^1^. Whereas the readout strategies for ChIP have evolved over 30 years from gel electrophoresis^1^ to massively parallel sequencing^2,3^, the fundamentals of ChIP have remained largely unchanged. Although ChIP-seq allows base-pair resolution mapping of TFs^4,5^, issues remain with high background that limits sensitivity, requirements for large number of cells, and artifacts resulting from cross-linking and solubilization^6-10^. Without an alternative method that is based on different principles from ChIP, it has been difficult to distinguish true positives from misleading false positive artifacts.

Alternative strategies have been used for the genome-wide mapping of protein-DNA interactions that can address some of these limitations of ChIP. For example, several methods, including DNase1 footpinting^11^, FAIRE-seq^12^, Sono-seq^13^, MNase-seq^14,15^ and ATAC-seq^16^, are being used to map TF binding genome-wide using a sequencing read-out. However, as these approaches are not targeted to specific proteins, they are not specific to any one TF. Furthermore they cannot be used to map specific chromatin states such as those demarcated by histone PTMs, which may be used to clinically differentiate healthy and disease states^17^.

Other methods provide target-specific mapping by genetically engineering a fusion between the protein of interest and an enzyme that methylates the surrounding DNA in the case of DamID^18^, or targeted cleavage of the protein’s footprint in the case of chromatin endogenous cleavage (ChEC)^19^. Enzyme tethering approaches are performed in vivo (DamID) or in situ (ChEC) without the need to fragment and solubilize chromatin. However, as they require a transgenic approach, this limits the scalability to large infrastructural consortiums such as ENCODE and the transferability to a clinical setting. In addition, these methods cannot map histone PTMs. These limitations were partially overcome by the chromatin immunocleavage (ChIC) method, whereby crude nuclei from crosslinked cells were first treated with a TF-specific antibody and then a fusion protein between protein A and Micrococcal Nuclease (pA-MN), which can be activated by calcium ions^19^. However, ChIC was developed using a Southern blot read-out, and so its applicability to genome-wide profiling remained unclear for over a decade.

We recently reported a major development of the ChIC strategy that we termed CUT&RUN (*c*leavage *u*nder *t*argets *& r*elease *u*sing *n*uclease; Fig. 1)^20^. Our protocol took unfixed nuclei and attached them to a solid support using concanavalin-A coated magnetic beads to allow simple handling. Following in situ binding of antibody and pA-MN specifically to the target protein, seconds after exposure to calcium at 0 °C, cleavage occurred on either side of the TF. As non-crosslinked nuclei were used, cleaved fragments released with two cuts were free to diffuse out of the nuclei, and so by simply pelleting the intact nuclei, the supernatant containing released chromatin fragments was used to extract DNA directly for sequencing. We found that performing the Ca^2+^-dependent digestion reaction at 0 °C was essential to limit the diffusion of the cleaved chromatin complexes, which would otherwise cleave and release accessible DNA. Overall, we showed that CUT&RUN has a much higher signal-to-noise ratio than crosslinking ChIP-seq, thereby allowing identification of previously unknown genomic features. CUT&RUN achieved base-pair resolution of mammalian TFs with only 10 million sequenced reads.

**Figure 1:**
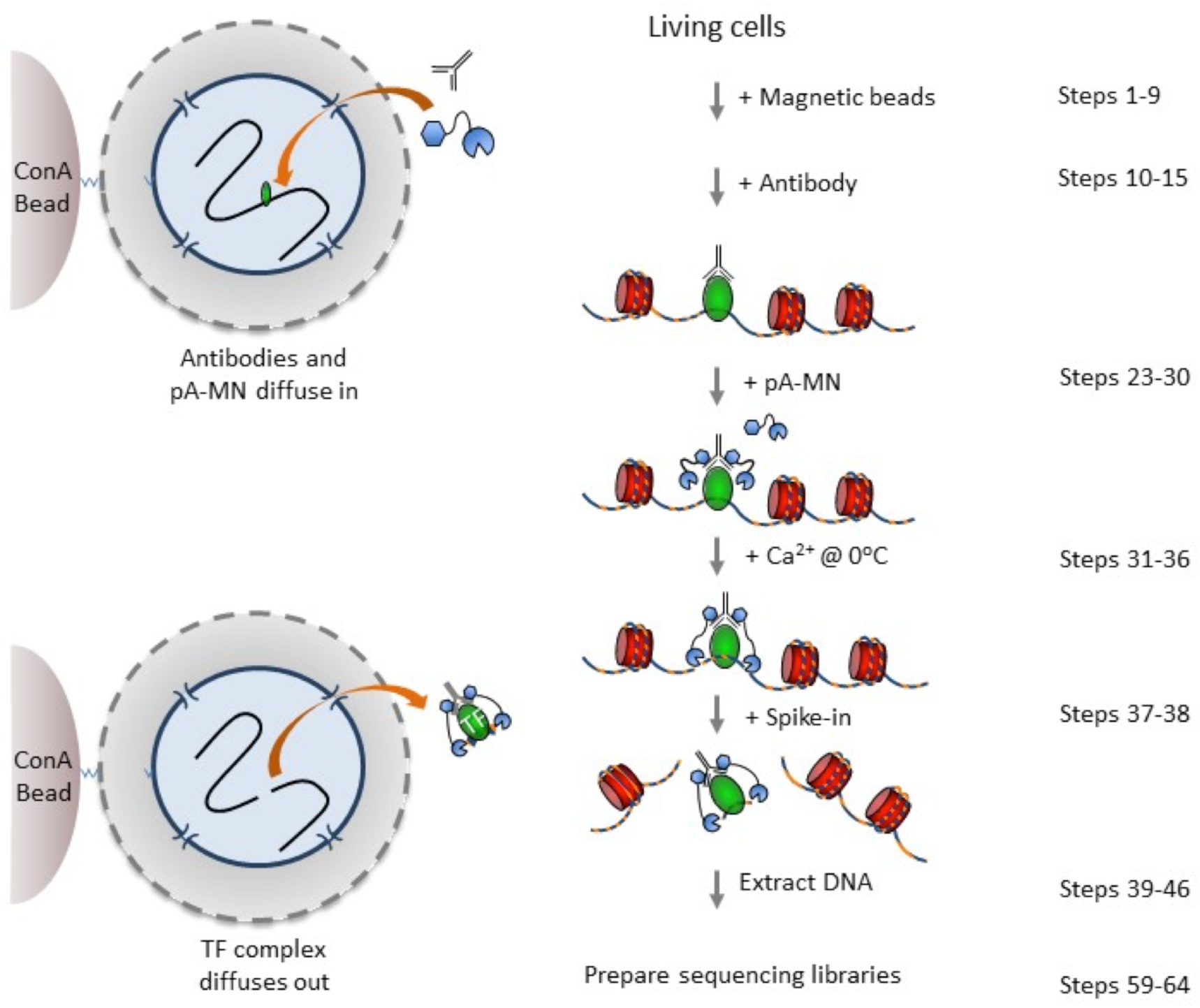
CUT&RUN requires less than a day from cells to DNA. A schematic overview of the CUT&RUN protocol. Cells are harvested and bound to concanavalin A-coated magnetic beads. Cell membranes are permeabilized with digitonin to allow the specific antibody to find it’s target. After incubation with antibody, beads are briefly washed, and then incubated with pA-MN. Cells are chilled to 0 ^°^C, and digestion begins with addition of Ca^2+^. Reactions are stopped by chelation including spike-in DNA and the DNA fragments released into solution by cleavage are extracted from the supernatant.

The need for quantitative mapping of protein-DNA interactions has become increasingly apparent^21^. However, due to the complexity of ChIP, which involves genome-wide solubilization of chromatin and immunoprecipitation, an involved quantitation strategy is required whereby a fixed number of cells from a different species that has antibody cross-reactivity is spiked-in^22^. The requirement for conserved epitopes limits general applicability. In contrast, due to the inherent simplicity of CUT&RUN, a straightforward spike-in strategy with heterologous DNA sufficed to accurately quantify binding events.

In summary, CUT&RUN has several advantages over ChIP-seq: (1) The method is performed in situ in non-crosslinked cells and does not require chromatin fragmentation or solubilization; (2) The intrinsically low background allows low sequence depth and identification of low signal genomic features invisible to ChIP; (3) The simple procedure can be completed within a day and is suitable for robotic automation; (4) The method can be used with low cell numbers compared to existing methodologies; (5) A simple spike-in strategy can be used for accurate quantitation of protein-DNA interactions. As such, CUT&RUN represents an attractive replacement for ChIP-seq, which is one of the most popular methods in biological research.

### Experimental Design

The CUT&RUN method for the in situ targeted cleavage and release of chromatin complexes is straightforward and can be completed in under a day using standard lab equipment. Here we provide a detailed protocol and also provide various options that might be used to tailor the protocol to specific situations. One of the strengths of CUT&RUN is that the entire reaction is performed in situ, whereby the antibody and pA-MN are free to diffuse into the nucleus. The original protocol used nuclei prepared by a combination of hypotonic lysis and treatment of cells with Triton X-100. This has been successful with a number of cell lines, but we have recently adapted the protocol to use cells permeabilized by the non-ionic detergent digitonin, which has been successfully used in other in situ methods, including ChEC-seq^23^ and ATAC-seq^24^. Digitonin partitions into membranes and extracts cholesterol. Membranes that lack cholesterol are minimally impacted by digitonin^25,26^. Nuclear envelopes are relatively devoid of cholesterol compared to plasma membranes. As such, treatment of cells with digitonin represents a robust method for permeabilizing cells without compromising nuclear integrity^26^. The protocol described here uses digitonin, but it is possible that individual experimental situations call for generating intact nuclei by other means, and such nuclei can be prepared by a suitable method, bound to concanavalin A-coated beads as per our previously published work and then enter the protocol below at step 10^20^.

One of the limitations of a protocol that has inherently low background and is amenable to low cell numbers is that the amount of DNA recovered can be very low, such that analysis even by sensitive capillary electrophoresis or picogreen assays (e.g. Agilent Tapestation and Qubit) are problematic. In addition, high resolution mapping techniques that cleave a minimal footprint are not suitable to PCR-based analysis of known binding loci, as it is not commonly possible to design ∼50 bp PCR amplicons. As such, we recommend using a positive control antibody that targets an abundant epitope and therefore the DNA can be readily detected. We have successfully used a rabbit monoclonal antibody raised against H3K27me3, with capillary electrophoresis showing with the amount of cleaved fragments being proportional to the number of starting cells. A nucleosomal ladder is expected by Tapestation or other sensitive electrophoretic analysis method (Fig. 2), and the use of a monoclonal antibody avoids potential lot-to-lot variation that can complicate troubleshooting. For less abundant epitopes such as CTCF, it is harder to detect the cleaved fragments by even sensitive electrophoretic analysis (Supplementary Figure 1). Once the expected digested DNA pattern is observed for the positive control by capillary electrophoresis such as H3K37me3, it is not necessary to sequence this sample. As a negative control, we recommend the use of a non-specific rabbit IgG antibody that will randomly coat the chromatin at low efficiency without sequence bias. We do not recommend a no-antibody control, as the lack of tethering increases the possibility that slight carry-over of pA-MN will result in preferential fragmentation of hyper-accessible DNA.

**Supplementary Figure S1:**
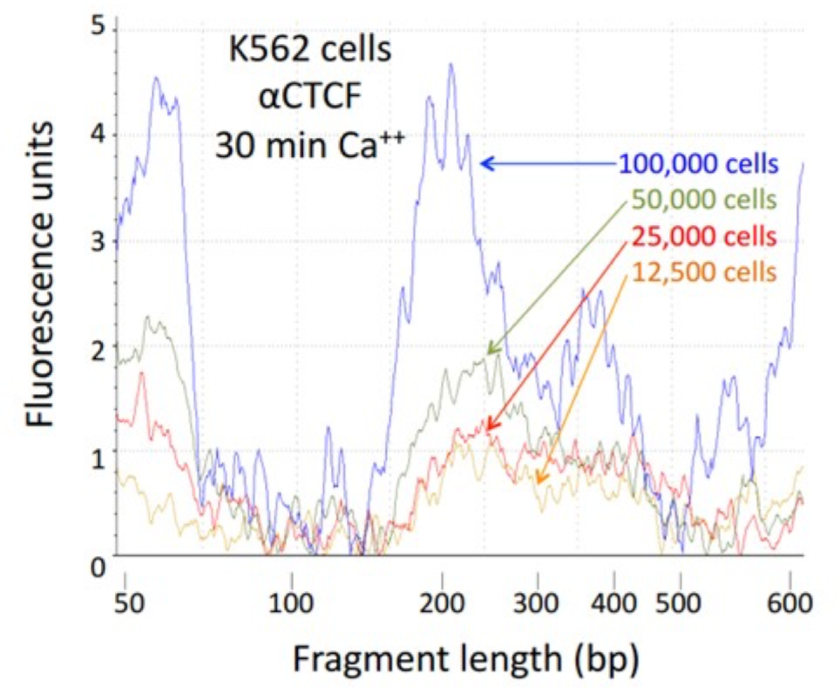
Tapestation analysis of CUT&RUN cleaved fragments using an anti-CTCF antibody. The remainder of these samples were used to make libraries for sequencing, with results shown in Figure 4.

**Supplementary Figure S2:**
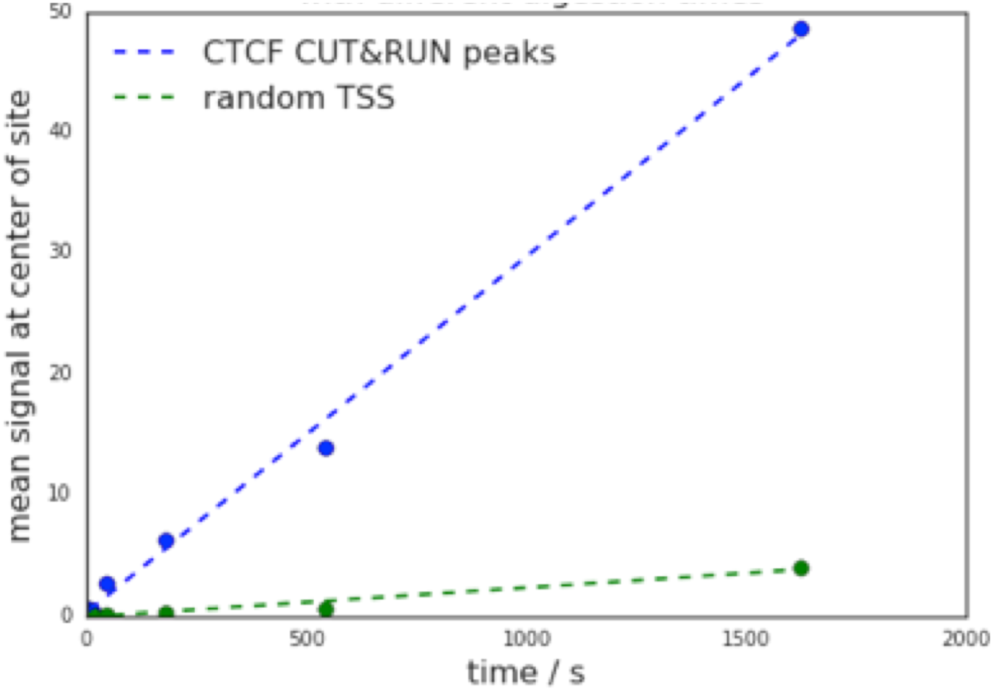
Yield increases with digestion time with little change in signal-to-noise. By scaling to spike-in DNA, quantitative measurement of the amount of cleaved DNA fragments is possible. The average signal over ∼20,000 CTCF CUT&RUN binding sites is compared to an equal number of non-overlapping transcriptional start sites (TSS) as a negative control regions. Spike-in scaled signal was summed over the −50 to +50 bp region relative to the center of the site or TSS.

**Figure 2:**
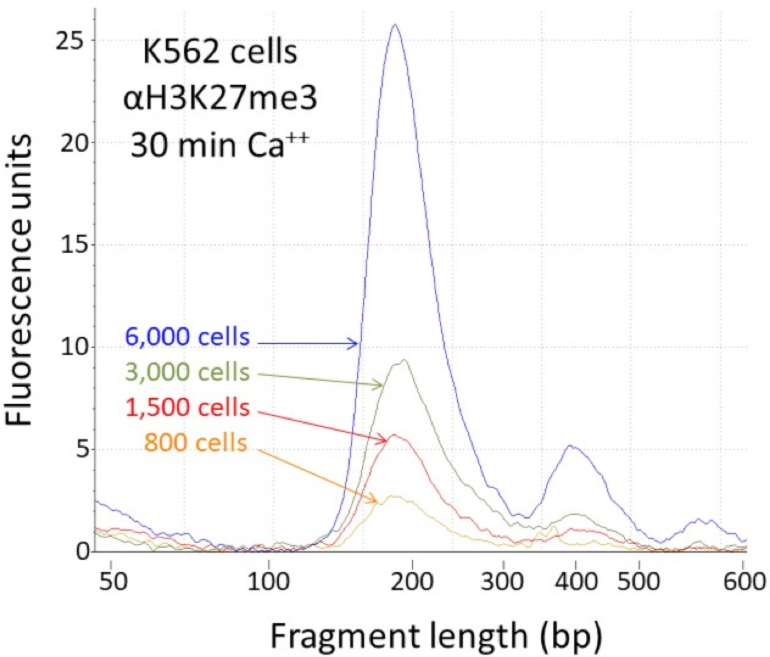
Tapestation analysis of an abundant histone epitope (H3K27me3) as a same-day positive control. The remainder of these samples were used to make libraries for sequencing, with results shown in Figure 3.

In our previously published study, we showed that targeted cleavage occurred within seconds of adding Ca^2+^ ions, and by virtue of being a sterically regulated tethered reaction, the cleavage pattern was constant over time. However, longer digestion times release more material with no apparent change in the signal-to-noise ratio (Supplementary Figure 2). We therefore recommend digesting for 30 minutes as a starting point that can be tailored based upon epitope abundance and antibody concentration.

### Applications of the method

CUT&RUN has the potential to replace all ChIP-based applications. For a typical research project in which ChIP-seq is currently used, transitioning to CUT&RUN is simple, as it can be done entirely on the benchtop using standard equipment that is already present in most molecular biology laboratories. Furthermore, as CUT&RUN is performed in situ in permeabilized cells that can readily be attached to a solid support such as magnetic beads, coated plates or glass slides, this method will readily transfer to robotics allowing high-throughput from cell to sequencing library. Adapting CUT&RUN to robotics should be more straightforward than is the case for ChIP-seq, as CUT&RUN does not require equipment such as sonicators or high speed spin steps to remove insoluble material that are difficult to automate.

Standard crosslinking ChIP protocols are not suitable for low cell numbers that are often obtained after fluorescence activated cell sorting or dissection, or in clinical settings. In light of this limitation, ATAC-seq has been used down to 5000 cells^24^. But ATAC-seq is limited to non-specific identification of TFs that are in accessible regions of chromatin and is unable to distinguish chromatin states demarcated by histone PTMs. Problems of epitope masking in crosslinking ChIP leading to low efficiency can be mitigated by using a native ChIP strategy, which was shown to provide high-quality data with as few as 5000 cells for abundant nucleosome epitopes, but was not applied to TFs^27^. Here, we show that CUT&RUN is suitable for application to 100 cells for profiling H3K27me3 or 1000 cells for CTCF sequence-specific DNA-binding protein. Therefore, CUT&RUN makes possible targeted genome-wide maps of protein-DNA interactions for rare cell types.

A recent advance in single-cell genomic analysis is *s*ingle-cell *c*ombinatorial *i*ndexing (“sci”), whereby split-pool barcoding is used to uniquely label a large number of intact individual cells without ever having to perform reactions on individual isolated cells. This approach has been successfully used for profiling transcriptomes^28^, chromatin accessibility (sci-ATAC-seq^29^), and 3-D interactions (sci-Hi-C^30^) in single cells. CUT&RUN, unlike ChIP, is performed inside intact permeabilized cells and therefore is amenable to combinatorial barcoding to map single-cell epitope-specific epigenomic landscapes.

Further development of the protocol could include a replacement for sequential ChIP to map co-occupancy of subunits within a protein complex. Sequential ChIP-seq has typically been challenging, and because of the very low yield after the second immunoprecipitation step, it is suitable only for abundant chromatin complexes. However, by first performing CUT&RUN, the cleaved chromatin complexes that are liberated into the supernatant at high efficiency could be immunoprecipitated with a second antibody. This application should allow compositional analysis and mapping of chromatin complexes genome-wide.

We previously showed that by virtue of CUT&RUN being an in situ cleavage approach and the inherent flexibility of the chromatin fiber, it is possible to probe the local chromatin structure including adjacent nucleosomes and 3D contacts. Hi-C, ChIA-PET and Hi-ChIP, which are popular technologies for genome-wide mapping of 3D nuclear organization, rely on formaldehyde crosslinking to stabilize protein-protein interactions^31-33^. As such, these techniques have no formal distance constraint for mapping a positive genomic interaction, as very large nuclear structures could be crosslinked. In contrast, TSA-seq^34^ and genome architecture mapping^35^ have distance constraints and therefore measure cytological distance, either by the limited diffusion of a reactive species or the cryosectioning of cells. Similarly, in CUT&RUN, the reach of protein A-MNase provides an intrinsic limit to how far cleavage can occur from an epitope and therefore how close two interacting DNA loci need to be in order to be cleaved by tethering to one of them. By combining CUT&RUN with a proximity based ligation method, it will be possible to generate factor-specific high resolution maps of nuclear architecture.

Other novel applications can be envisioned. Any epitope for which an antibody is available can potentially be subjected to profiling using CUT&RUN, and CUT&RUN in situ mapping of lncRNAs would seem to be an attractive alternative to DRIP-seq^36^. In addition, the ability of CUT&RUN to profile insoluble chromatin^20^ suggests that combining CUT&RUN with salt fractionation will allow for an epigenomic map to be based on chromatin solubility, which has traditionally been used to define classical “active” chromatin^37-39^. In this way, each DNA-binding protein or chromatin feature being profiled can be enriched with information about its solubility, a key physical property. Although salt-fractionation can be performed with MNase-based ChIP-seq^39^ high salt can disrupt the complex and cause loss of the epitope prior to antibody binding, whereas with CUT&RUN, salt fractionation is performed only after the antibody is bound and the fragments cleaved.

### Comparison with other methods

Table 1 lists metrics for CUT&RUN and three ChIP-seq methods, X-ChIP-seq^3^, ChIP-exo^4^ and N-ChIP-seq^40^. Compared to these ChIP-seq methods, CUT&RUN requires fewer cells and fewer reads, has a higher signal-to-noise ratio, has no fragmentation bias, is faster and is amenable to spike-in for quantitation.

**Table 1:**
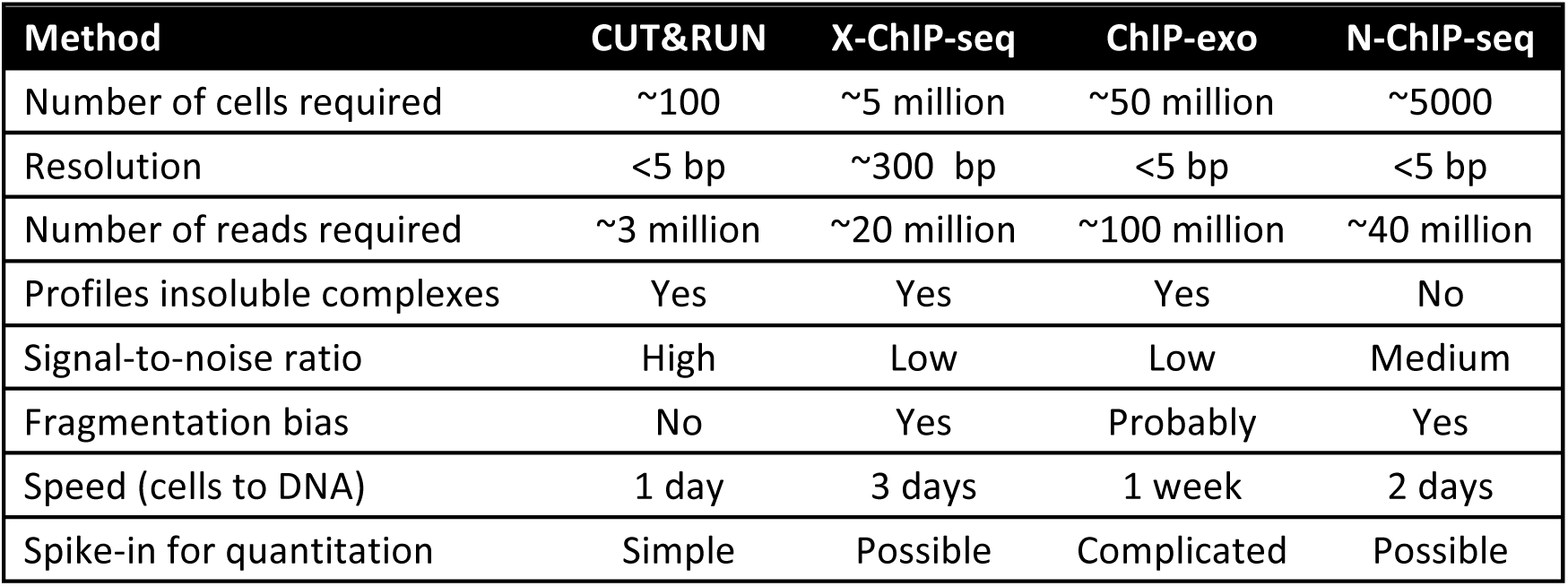
Comparison of CUT&RUN to ChIP-seq protocols.

| Method | CUT&RUN | X-ChIP-seq | ChIP-exo | N-ChIP-seq |
| --- | --- | --- | --- | --- |
| Number of cells required | ~100 | ~5 million | ~50 million | ~5000 |
| Resolution | <5 bp | ~300 bp | <5 bp | <5 bp |
| Number of reads required | ~3 million | ~20 million | ~100 million | ~40 million |
| Profiles insoluble complexes | Yes | Yes | Yes | No |
| Signal-to-noise ratio | High | Low | Low | Medium |
| Fragmentation bias | No | Yes | Probably | Yes |
| Speed (cells to DNA) | 1 day | 3 days | 1 week | 2 days |
| Spike-in for quantitation | Simple | Possible | Complicated | Possible |

An important advance in ChIP-based technologies has been to leverage next generation sequencing to generate base-pair resolution genome-wide maps of protein-DNA interactions^41^.

In contrast to standard crosslinking ChIP where sonication is used to fragment the chromatin to a minimum of ∼200 bp fragments, exonuclease treatment in ChIP-exo or MNase digestion in high-resolution X-ChIP-seq or native ChIP approaches allows limit or near-limit digestion^4,5,20,40,42^. However, this improvement in resolution in crosslinking strategies has often come at the price of increases in sequence depth requirements and the number of cells required. For example, in ChIP-exo, any sonicated fragments that contain more than just the target protein, such as an adjacent nucleosome, will form a block to the exo-nuclease in generating minimal TF footprints and as such contribute to an apparent localized background, requiring increased cell numbers and sequencing depths to call high resolution peak pairs. Native ChIP often does not suffer from these associated problems, but has limited general applicability due to the requirement to generate soluble chromatin extracts in the absence of harsh detergents and therefore is best suited to stably bound proteins and may require optimization on a case-by-case basis. It has previously been shown that sonication, such as is used for cross-linking ChIP methods, is non-random and therefore is subject to a fragmentation bias^5,43^. As CUT&RUN is performed on intact cells or nuclei without fragmentation, it can be used to probe all genomic compartments. Technologies that use MNase for genome-wide digestion can suffer from A/T bias of the enzyme^44^ and will preferentially digest open chromatin. In contrast, CUT&RUN involves a sterically regulated cleavage reaction, and we have shown that it does not suffer from any detectable A/T or DNA accessibility bias^20^.

### Limitations

As is the case with ChIP, the success of CUT&RUN depends in large part on the affinity of the antibody for its target and its specificity under the conditions used for binding. Because antibodies bind to their epitopes in the solid state using CUT&RUN, we would expect that antibodies successfully tested for specificity by immunofluorescence (IF) would be likely to work with CUT&RUN, with the caveat that IF generally involves fixation, whereas formaldehyde fixation decreases the efficiency of CUT&RUN.

In the standard CUT&RUN protocol we recommend allowing the cleaved chromatin complexes to diffuse out of the nuclei, thereby permitting simple isolation of the cut DNA from the supernatant fraction with the undigested genome retained in the intact nuclei. However, it is possible that a chromatin complex is too large to diffuse out or that protein-protein interactions retain the cleaved complex. In such cases, total DNA may be extracted after the digestion. By doing a very simple size selection using ½ volume of paramagnetic carboxylated beads (e.g. Agencourt AMPure XP beads) fragments below ∼700 bp will be selected for. We previously showed that this strategy was successful for the ∼1 MDa yeast RSC complex^20^.

## MATERIALS

### REAGENTS

- Cell suspension. We have used human K562 cells, Drosophila S2 cells and dissected Drosophila tissues such as brains and imaginal disks, and spheroplasted yeast.
- Concanavalin-coated magnetic beads (Bangs Laboratories, ca. no. BP531)
- Antibody to an epitope of interest. For example, rabbit α-CTCF polyclonal antibody (Millipore 07-729) for mapping 1D and 3D interactions by CUT&RUN
- Positive control antibody to an abundant epitope, *e.g.* α-H3K27me3 rabbit monoclonal antibody (Cell Signaling Technology, cat. no. 9733)
- Negative control antibody to an absent epitope, *e.g.* guinea pig α-rabbit antibody
- 5% Digitonin (EMD Millipore, cat. no. 300410)
- Protein A–Micrococcal Nuclease (pA-MNase) fusion protein (provided in 50% glycerol by the authors upon request). Store at −20 ^°^C.
- Spike-in DNA (e.g., from *Saccharomyces cerevisiae* micrococcal nuclease-treated chromatin, provided by authors upon request)
- Distilled, deionized or RNAse-free H_2_O (dH_2_O e.g., Promega, cat. no. P1197)
- 1 M Manganese Chloride (MnCl_2_; Sigma-Aldrich, cat. no. 203734)
- 1 M Calcium Chloride (CaCl_2_; Fisher, cat. no. BP510)
- 1 M Potassium Chloride (KCl; Sigma-Aldrich, cat. no. P3911)
- 1 M Hydroxyethyl piperazineethanesulfonic acid pH 7.5 (HEPES (Na^+^); Sigma-Aldrich, cat. no. H3375)
- 1 M Hydroxyethyl piperazineethanesulfonic acid pH 7.9 (HEPES (K^+^); Sigma-Aldrich, cat. no. H3375)
- 5 M Sodium chloride (NaCl; Sigma-Aldrich, cat. no. S5150-1L)
- 0.5 M Ethylenediaminetetraacetic acid (EDTA; Research Organics, cat. no. 3002E)
- 0.2 M Ethylene glycol-bis(β-aminoethyl ether)-N,N,N’,N’-tetraacetic acid (EGTA; Sigma-Aldrich, cat. no. E3889)
- 2 M Spermidine (Sigma-Aldrich, cat. no. S2501)
- Roche Complete Protease Inhibitor EDTA-Free tablets (Sigma-Aldrich, cat. no. 5056489001)
- 2 mg/ml Glycogen (1:10 dilution of Sigma-Aldrich, cat. no. 10930193001)
- RNase A, DNase and protease-free (10 mg/ml; Thermo Fisher Scientific, cat. no. EN0531)
- Gel and PCR Clean-up kit (Macherey-Nagel NucleoSpin^®^, cat. no. 740609.250)
- Agencourt AMPure XP magnetic beads (Beckman Coulter, cat. no. A63880)
- 10% Sodium dodecyl sulfate (SDS; Sigma-Aldrich, cat. no. L4509)
- Proteinase K (Thermo Fisher Scientific, cat. no. EO0492)
- Phenol-chloroform-isoamyl alcohol 25:24:1 (PCI; Invitrogen, cat. no. 15593049)
- Chloroform (Sigma, cat. no. 366919-1L)
- 1 M Tris-HCl pH 8.0
- Ethanol (Decon Labs, cat. no. 2716)
- Qubit dsDNA HS kit (Life Technologies, cat. no. Q32851)

### EQUIPMENT

- Centrifuge Eppendorf 5810, swinging bucket
- Centrifuge Eppendorf 5424, fixed angle rotor
- Centrifuge Eppendorf 5415R, refrigerated fixed angle rotor
- Macsimag magnetic separator (Miltenyi, cat. no. 130-092-168), which allows clean withdrawal of the liquid from the bottom of 1.7 and 2 ml microfuge tubes.
- Vortex mixer (e.g., VWR Vortex Genie)
- Micro-centrifuge (e.g., VWR Model V)
- 1.5-ml microcentrifuge tubes (Genesee, cat. no. 22-282)
- 2-ml microcentrifuge tubes (Axygen, cat. no. MCT-200-C)
- Tube rotator (Labquake, Thermo Fisher)
- Heater block with wells for 1.5-ml microcentrifuge tubes
- Water baths (set to 37 °C and 70 °C)
- MaXtract phase-lock microcentrifuge tubes (Qiagen, cat. no. 139046)
- Capillary electrophoresis instrument (e.g. Agilent Tapestation 4200)
- Qubit Fluorometer (Life Technologies, cat. no. Q33216)

### REAGENT SETUP

**5% Digitonin** To reconstitute enough digitonin for an experiment, weigh out the powder in a 2 ml microcentrifuge tube, boil water in a small beaker in a microwave oven, and pipette in and out to warm the 1000 μL pipette tip. Pipette the hot water into the tube with the digitonin powder to make 5% (w/v), close the cap and quickly vortex on full until the digitonin is completely dissolved. If refrigerated, this stock can be used within a week, but will need reheating as the digitonin slowly precipitates. The effectiveness of digitonin varies between batches, so testing permeability of Trypan blue is recommended to determine the concentration to use for a cell type. We have obtained excellent results for K562 cells with 0.02-0.1% digitonin.

- **CAUTION:** Digitonin is toxic and care should be taken especially when weighing out the powder. A digitonin stock may be prepared by dissolving in dimethylsulfoxide (DMSO), but be aware that DMSO can absorb through the skin.

**Binding buffer** Mix 400 μL 1M HEPES-KOH pH 7.9, 200 μL 1M KCl, 20 μL 1M CaCl_2_ and 20 μL 1M MnCl_2_, and bring the final volume to 20 ml with dH_2_O. Store the buffer at 4 ^°^C for 6 months.

**Concanavalin A-coated beads** Gently resuspend and withdraw enough of the slurry such that there will be 10 μL for each final sample and/or digestion time point. Transfer into 1.5 ml Binding buffer in a 2 ml tube. Place the tube on a magnet stand to clear (30 s to 2 min). Withdraw the liquid, and remove from the magnet stand. Add 1.5 ml Binding buffer, mix by inversion or gentle pipetting, remove liquid from the cap and side with a quick pulse on a micro-centrifuge. Resuspend in a volume of Binding buffer equal to the volume of bead slurry (10 μL per final sample).

**Wash buffer** Mix 1 ml 1 M HEPES pH 7.5, 1.5 ml 5 M NaCl, 12.5 μL 2 M Spermidine, bring the final volume to 50 ml with dH_2_O, and add 1 Roche Complete Protease Inhibitor EDTA-Free tablet. Store the buffer at at 4 ^°^C for up to 1 week.

**Dig-wash buffer** Mix 160-800 μL 5% Digitonin with 40 ml Wash buffer. The effectiveness of digitonin varies between batches, so testing permeability of Trypan blue is recommended to determine the concentration to use. We have obtained excellent results for K562 cells with 0.02-0.1% digitonin. Store the buffer at at 4 ^°^C for up to 1 day.

**Antibody buffer** Mix 8 μL 0.5 M EDTA with 2 ml Dig-wash buffer and place on ice. Divide into aliquots for each antibody and add antibody solution or serum to a final concentration of 1:100 or to the manufacturer’s recommended concentration for immunofluorescence.

**2XSTOP** To 4.2 ml dH_2_O add 340 μl 5M NaCl, 200 μL 0.5M EDTA, 100 μL 0.2M EGTA, 20 μL 5% Digitonin, 25 μL RNase A, 125 μL 2 mg/ml glycogen and 2 pg/ml heterologous spike-in DNA. Store the buffer at 4 ^°^C for up to 1 week.

- **CRITICAL STEP:** Heterologous spike-in DNA for calibration should be fragmented down to ∼200 bp mean size, for example an MNase-treated sample of mononucleosome-sized fragments. As we use the total number of mapped reads as a normalization factor only, very little spike-in DNA is needed. For example, addition of 1.5 pg results in 1,000-10,000 mapped spike-in reads for 1-10 million mapped experimental reads (in inverse proportion).

## PROCEDURE

### Binding cells to beads

- **TIMING 30 min**
- **CRITICAL STEP:** All steps prior to the addition of antibody are performed at room temperature to minimize stress on the cells. Because it is crucial that DNA breakage is minimized throughout the protocol, we recommend that cavitation during resuspension and vigorous vortexing be avoided. 1) Harvest fresh culture(s) at room temperature and count cells. The same protocol can be used for 100 to 250,000 mammalian cells per sample and/or digestion time point.
- **PAUSE POINT:** If necessary, cells can be cryopreserved in 10% DMSO using a Mr. Frosty isopropyl alcohol chamber. We do not recommend flash freezing, as this can cause background DNA breakage that may impact final data quality. 2) Centrifuge 3 min 600 × g at room temperature and withdraw liquid. 3) Resuspend in 1.5 ml room temperature Wash buffer by gently pipetting and transfer if necessary to a 2 ml tube. 4) Centrifuge 3 min 600 × g at room temperature and withdraw liquid. 5) Repeat steps 3 and 4. 6) Resuspend in 1 ml room temperature Wash buffer by gently pipetting. 7) While gently vortexing the cells at room temperature, add the bead slurry. 8) Rotate 5-10 min at room temperature. 9) Divide into aliquots in 1.5-ml tubes, one for each antibody to be used.
- **CRITICAL STEP:** To evaluate success of the procedure without requiring library preparation, include in parallel a positive control antibody (*e.g.* α-H3K27me3) and a negative control antibody (*e.g.* α-rabbit). Do not include a no-antibody control, as the lack of tethering may allow any unbound pA-MN to act as a “time-bomb” and digest accessible DNA, resulting in a background of DNA-accessible sites.

### Bind (primary) antibodies

- **TIMING 15 min–overnight, with longer incubations providing higher yields** 10) Place on the magnet stand to clear and pull off the liquid.
- **CRITICAL STEP:** Although low-retention pipette tips are preferred for accurate solution transfers, use only conventional (not low-binding) microcentrifuge tubes to avoid loss of beads while decanting. 11) Place each tube at a low angle on the vortex mixer set to low (∼1100 rpm) and squirt 50 μL of the Antibody buffer (per sample and/or digestion time point) along the side while gently vortexing to allow the solution to dislodge most or all of the beads. Tap to dislodge the remaining beads.
- **CRITICAL STEP:** The presence of EDTA during antibody treatment removes excess divalent cation used to activate the ConA, because carry-over of Ca++ from the beads can prematurely initiate strand cleavage after addition of pA-MN. Chelation of divalent cations when cells are permeabilized also serves to quickly halt metabolic processes and prevent endogenous DNAse activity. Washing out the EDTA before pA-MN addition avoids inactivating the enzyme. Spermidine in the wash buffer is intended to compensate for removal of Mg++, which might otherwise affect chromatin properties. 12) Place on the tube rotator at 4 ^°^C for ∼2 hr, or at room temperature for 5-10 min.
- **PAUSE POINT** Antibody incubation may proceed overnight at 4 ^°^C. 13) Remove liquid from the cap and side with a quick pulse on a micro-centrifuge.
- **CRITICAL STEP:** After mixing, but before placing a tube on the magnet stand, a very quick spin on a micro-centrifuge (no more than 100 × g) will minimize carry-over of antibody and pA-MN that could result in overall background cleavages during the digestion step. 14) Place on the magnet stand to clear (∼30 s) and pull off all of the liquid. 15) Add 1 ml Dig-wash buffer, mix by inversion, or by gentle pipetting using a 1 ml tip if clumps persist, and remove liquid from the cap and side with a quick pulse on a micro-centrifuge.

### Bind secondary antibody (as required)

- **TIMING 15 min-1.5 hr**
- **CRITICAL STEP:** The binding efficiency of Protein A to the primary antibody depends on host species and IgG isotype. For example, Protein A binds well to rabbit and guinea pig IgG but poorly to mouse and goat IgG, and so for these latter antibodies a secondary antibody, such as rabbit α-mouse is recommended. 16) Place on the magnet stand to clear and pull off all of the liquid. 17) Place each tube at a low angle on the vortex mixer set to low (∼1100 rpm) and squirt 50 μL of the Dig-wash buffer (per sample and/or digestion time point) along the side while gently vortexing to allow the solution to dislodge most or all of the beads. Tap to dislodge the remaining beads. 18) Mix in the secondary antibody to a final concentration of 1:100 or to the manufacturer’s recommended concentration for immunofluorescence. 19) Place on the tube rotator at 4 ^°^C for ∼1 hr, or at room temperature for 5-10 min. 20) Remove liquid from the cap and side with a quick pulse on a micro-centrifuge. 21) Place on the magnet stand to clear and pull off all of the liquid. 22) Add 1 ml Dig-Wash buffer, mix by inversion, or by gentle pipetting if clumps persist, and remove liquid from the cap and side with a quick pulse on a micro-centrifuge.

### Bind Protein A-MNase fusion protein

- **TIMING 15 min–1.5 hr** 23) Place on the magnet stand to clear and pull off all of the liquid. 24) Place each tube at a low angle on the vortex mixer set to low (∼1100 rpm) and squirt 50 μL of the Dig-wash buffer (per sample and/or digestion time point) along the side while gently vortexing to allow the solution to dislodge most or all of the beads. Tap to dislodge the remaining beads. 25) Mix in the pA-MNase to a final concentration of ∼700 ng/ml (e.g. 2.5 μL/50 μL of a 1:10 dilution of the 140 μg/ml glycerol stock provided upon request). 26) Place on the tube rotator at 4 ^°^C for ∼1 hr, or at room temperature for 5-10 min. 27) Remove liquid from the cap and side with a quick pulse on a micro-centrifuge. 28) Place on the magnet stand to clear and pull off all of the liquid. 29) Add 1 ml Dig-wash buffer, mix by inversion, or by gentle pipetting if clumps persist, and remove liquid from the cap and side with a quick pulse on a micro-centrifuge. 30) Repeat Dig-wash steps 28-29.

### Targeted digestion

- **TIMING 45 min** 31) Place on the magnet stand to clear and pull off all of the liquid. 32) Place each tube at a low angle on the vortex mixer set to low (∼1100 rpm) and add 100 μL of the Dig-wash buffer (per sample and/or digestion time point) along the side while gently vortexing to allow the solution to dislodge most or all of the beads. Tap to dislodge the remaining beads. 33) Insert tubes into the 1.5 ml wells of a heater block sitting in wet ice to chill down to 0 ^°^C. 34) Remove each tube from the block, mix in 2 μL 100 mM CaCl_2_ (diluted 1:10 from a 1 M stock) with gentle vortexing and immediately replace the tube in the 0 ^°^C block. 35) Incubate at 0 ^°^C for the desired digestion time (default is 30 min).
- **CRITICAL STEP:** MNase binds DNA but only cleaves when Ca^++^ is present, so that digestion is a zero-order reaction that seems to be less temperature-dependent than the subsequent diffusion of released pA-MNase-bound particles that can digest accessible regions of the genome. Cleavage and release of particles in most of the cell population can be obtained at 0 ^°^C while minimizing background cleavages attributable to diffusion. We have found that digestion at ambient temperature or higher results in unacceptable background cleavage levels. 36) Add 100 μL 2XSTOP and mix by gentle vortexing. When there are multiple time points, remove 100 μL to 100 μL 2XSTOP and mix by gentle vortexing.
- **CRITICAL STEP:** Heterologous spike-in DNA should be present in the 2XSTOP to calibrate DNA amounts, for example to compare treatments or digestion time points. This is especially important for CUT&RUN, as there is too little background cleavage for normalization of samples.

### Target chromatin release

- **TIMING 20 min** 37) Incubate 10 min 37 ^°^C to release CUT&RUN fragments from the insoluble nuclear chromatin. 38) Centrifuge 5 min 4 ^°^C 16,000 × g and place on magnet stand.

### Option A: Fast DNA extraction by spin column

- **TIMING 20 min** 39) Place a spin column into a collection tube and add 400 μL Buffer NT1 (from NucleoSpin kit or equivalent). 40) Decant the supernatant cleanly from the pellet and transfer to the NT1 in the spin column pipetting gently up and down to mix. 41) Centrifuge 30 s at 11,000 × g. Discard flow-through. 42) Add 700 μL Buffer NT3. Centrifuge 30 s 11,000 × g. Discard flow-through. 43) Add 700 μL Buffer NT3. Centrifuge 30 s @11,000 × g. Discard flow-through and replace in rotor. 44) Centrifuge for 1 min 11,000 × g. Let dry 5 min. 45) Place in a fresh tube and add 20-40 μL Buffer NE to membrane. 46) After 1 min, centrifuge for 1 min @11,000 × g.

### Option B: Alternate DNA extraction (preferred for quantitative recovery of ≤80 bp fragments)

- **TIMING 1.5 hr** 47) Decant the supernatant cleanly from the pellet and transfer to a fresh 1.5-ml microcentrifuge tube. 48) To each sample add 2 μL 10% SDS (to 0.1%), and 2.5 μL Proteinase K (20 mg/ml). Mix by inversion and incubate 10 min 70 ^°^C. 49) Add 300 μL PCI and mix by full-speed vortexing ∼2 s. 50) Transfer to a phase-lock tube, and centrifuge 5 min room temperature at 16,000 × g. 51) Add 300 μL chloroform and invert ∼10x to mix. 52) Remove liquid by pipetting to a fresh tube containing 2 μL 2 mg/ml glycogen. 53) Add 750 μL 100% ethanol and mix by vortexing or tube inversion. 54) Chill on ice and centrifuge 10 min at 4 °C 16,000 × g. 55) Pour off the liquid and drain on a paper towel. 56) Rinse the pellet in 1 ml 100% ethanol and centrifuge 1 min at 4 °C 16,000 × g. 57) Carefully pour off the liquid and drain on a paper towel. Air dry. 58) When the pellet is dry, dissolve in 25-50 μL 1 mM Tris-HCl pH8 0.1 mM EDTA.

### Library preparation and sequencing

- **TIMING2–4 d** 59) Optional: Quantify 1-2 μL, for example using fluorescence detection with a Qubit instrument. 60) Optional: Evaluate the presence of cleaved fragments and the size distribution by capillary electrophoresis with fluorescence detection, for example using a Tapestation instrument.
- **CRITICAL STEP:** Some long undigested DNA will leak through, and this is what will dominate the Qubit fluorescence for CUT&RUN of typical transcription factors. For these, the targeted DNA recovered is too low in amount and too small in size to be detected by gel analysis or even by Tapestation. In such cases it may be necessary to make a PCR-amplified library to quantify by Tapestation or Bioanalyzer analysis. 61) Prepare barcoded libraries for Illumina sequencing with Tru-Seq adapters using a single-tube protocol, following the manufacturer’s instructions. Rapid PCR cycles favor exponential amplification of the desired CUT&RUN fragments over linear amplification of large DNA fragments that are too long for polymerase to complete.
- **CRITICAL STEP:** To minimize the contribution of large DNA fragments, PCR cycles should be at least 12-14 cycles, preferably with a 10 s 60^°^C combined annealing/extension step. Good results have been obtained with the Hyper-prep kit (KAPA Biosystems). 62) Quantify library yield using dsDNA-specific assay, such as Qubit. 63) Determine the size distribution of libraries by Agilent 4200 TapeStation analysis. 64) Perform paired-end Illumina sequencing on the barcoded libraries following the manufacturer’s instructions.
- **CRITICAL STEP:** Because of the very low background with CUT&RUN, typically 5 million paired-end reads suffices for transcription factors or nucleosome modifications, even for the human genome. For maximum economy, we mix up to 24 barcoded samples per lane on a 2-lane flow cell, and perform paired-end 25x25 bp sequencing. Single-end sequencing is not recommended for CUT&RUN, as it sacrifices resolution and discrimination between transcription factors and neighboring nucleosomes.

### Data processing and analysis

- **TIMING1 d (variable)** 65) We align paired-end reads using Bowtie2 version 2.2.5 with options: --local --very-sensitive-local --no-unal --no-mixed --no-discordant --phred33 −I 10 −X 700. For mapping spike-in fragments, we also use the --no-overlap --no-dovetail options to avoid cross-mapping of the experimental genome to that of the spike-in DNA.
- **CRITICAL STEP:** Separation of sequenced fragments into ≤120 bp and ≥150 bp size classes provides mapping of the local vicinity of a DNA-binding protein, but this can vary depending on the steric access to the DNA by the tethered MNase. Single-end sequencing is not recommended for CUT&RUN, as it sacrifices resolution and discrimination between transcription factors and neighboring nucleosomes. 66) Scripts available from https://github.com/peteskene are customized for processing, spike-in calibration, and analysis CUT&RUN data.

## TROUBLESHOOTING

**Table 2:** Troubleshooting table.

| Steps | Problem | Possible reasons | Solutions |
| --- | --- | --- | --- |
| 13 | Beads clump and cannot be disaggregated | Cells lyse | Reduce the digitonin concentration |
| 59 | No DNA is detected by Qubit fluorimetry | This is typical for low cell numbers (<10,000 cells) but otherwise may indicate an antibody failure. | Replace antibody. Antibody binding may be tested by immunofluorescence. |
| 60 | No DNA <200 bp is detected by Tapestation analysis | This is typical for most DNA-binding proteins, but otherwise may indicate failure of antibody binding or digestion. | Run a positive control sample for an abundant epitope, *e.g.* H3K27me3. |
| 60 | A nucleosome ladder is detected by Tapestation analysis | This is typical for abundant nucleosomal epitopes, but otherwise may indicate release of pA-MNase during digestion. | Run negative control sample using an IgG, *e.g.* guinea pig α-rabbit. |
| 60 | Small DNA or a ladder is seen in the negative control by Tapestation analysis | Divalent cations have not been removed by the EDTA in the antibody solution, or the negative control antibody failed to bind, allowing the pA-MNase to behave as a “time-bomb” when $\text{Ca}^{++}$ is added. | Replace antibody. Reduce pA-MN concentration. Reduce digestion time. Add a third wash step before digestion. |

## TIMING

### Day 1 Cells to DNA

Steps 1-9 30 min

Steps 10-15 2.5 hr

Steps 16-22 (Optional) 1.5 hr

Steps 23-30 1.5 hr

Steps 31-36 1 hr

Steps 37-38 20 min

Steps 39-46 20 min

Steps 47-58 (Alternative to Steps 39-46) 1.5 hr

**Days 2-4 Library preparation and sequencing**

Steps 59-64

**Day 5 (variable) Data processing and analysis**

Steps 65-66

## ANTICIPATED RESULTS

Human K562 cells were cultured at 37 ^°^C, counted, harvested at 1 × 10^6^ cells/ml by low-speed centrifugation, suspended and pelleted twice in Wash buffer, then diluted and mixed with wash buffer in a 300 μL volume to achieve a doubling series between 50 and 6,000 cells. Ten μL of a Ca^2+^- and Mn^2+^-washed ConA-coated magnetic bead slurry was added in binding buffer to each cell suspension with gentle vortexing. After 10 min, cells were collected on a magnet stand, decanted, resuspended in 50 μL Antibody buffer containing anti-H3K27me3 (1:100, CST #9733), 2 mM EDTA and 0.05% digitonin and incubated at 4 ^°^C for 15 hr. After collecting the beads on a magnet stand and washing once in 1 ml cold Dig-wash, cells were resuspended in 100 μL pA-MN (1:500 360 μg/ml) in Dig-wash and incubated at 4 ^°^C for 1 hr. Beads were collected on a magnet stand, washed twice in 1 ml Dig-wash, resuspended in 150 μL Dig-wash, and chilled to 0 ^°^C. Three μL 100 mM CaCl_2_ was added, and 0 oC incubation was continued for 30 min. Reactions were terminated with 1 vol 2XSTOP, incubated at 37 ^°^C for 20’ and centrifuged at 4 ^°^C for 5’ 16,000 xg. Both the supernatant and pellet were extracted following Steps 47-58). DNA from pellets was quantified by Qubit fluorescence (Supplementary Figure 3). DNA from selected supernatant fractions was resolved by Tapestation analysis (Fig. 2) and subjected to Illumina PE25x25 sequencing.

**Supplementary Figure S3:**
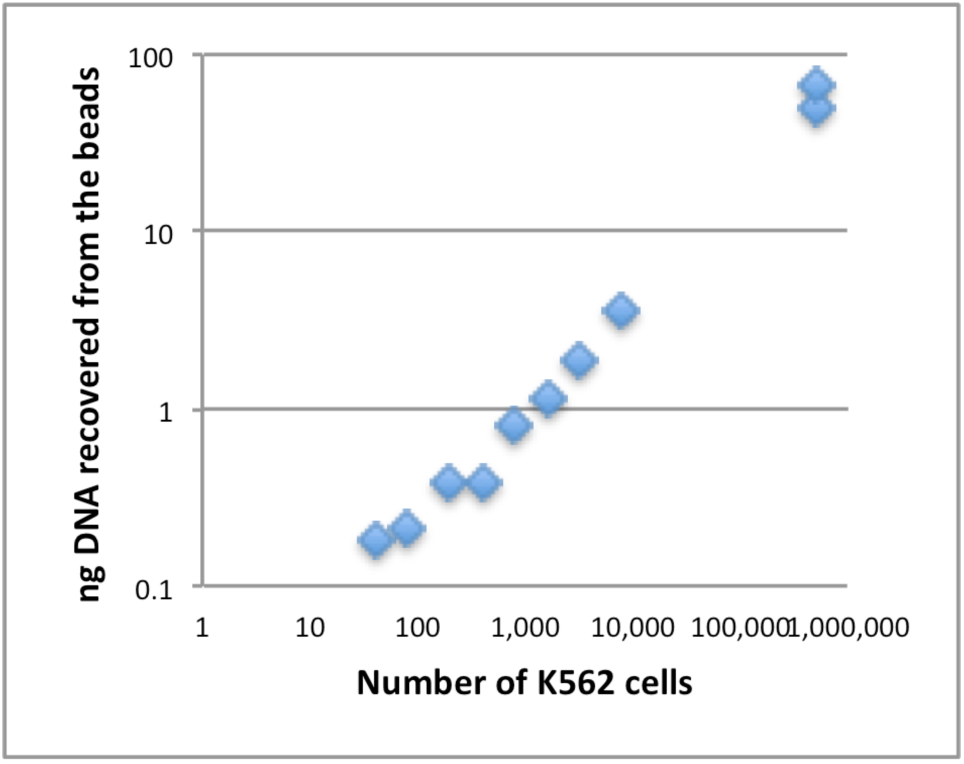
Recovery of insoluble DNA after release of H3K27me3 DNA into the supernatant. DNA was extracted from bead pellets of the samples used in the experiment described in Fig. 2.

Typical ChIP-seq experiments use high starting cell numbers that result in a large number of unique sonicated fragments that are immunoprecipitated. In contrast, as CUT&RUN allows low cell numbers and has a relatively low background, the number of unique fragments is less than typical sequence depths. Therefore, high sequencing depths from low cell number experiments could result in redundant sequencing of PCR duplicates. Presumed PCR duplicates were removed and mapped fragments were randomly sampled without replacement, resulting in 7.5 million unique reads per sample, displayed as normalized counts from stacked reads (Fig. 3). For comparison, a sample of 7.5 million unique reads were sampled from an ENCODE dataset for H3K27me3 in K562 cells. It is evident that very little loss of data quality occurred with reduction in cell number down to 100 cells. In contrast, the ENCODE profile sampled at the same depth shows a blurry profile owing to the high background inherent to ChIP. CUT&RUN using an anti-CTCF antibody (1:100, Millipore 07-729) was performed similarly, yielding profiles with little loss of data quality down to 1000 cells (Fig. 4).

**Figure 3:**
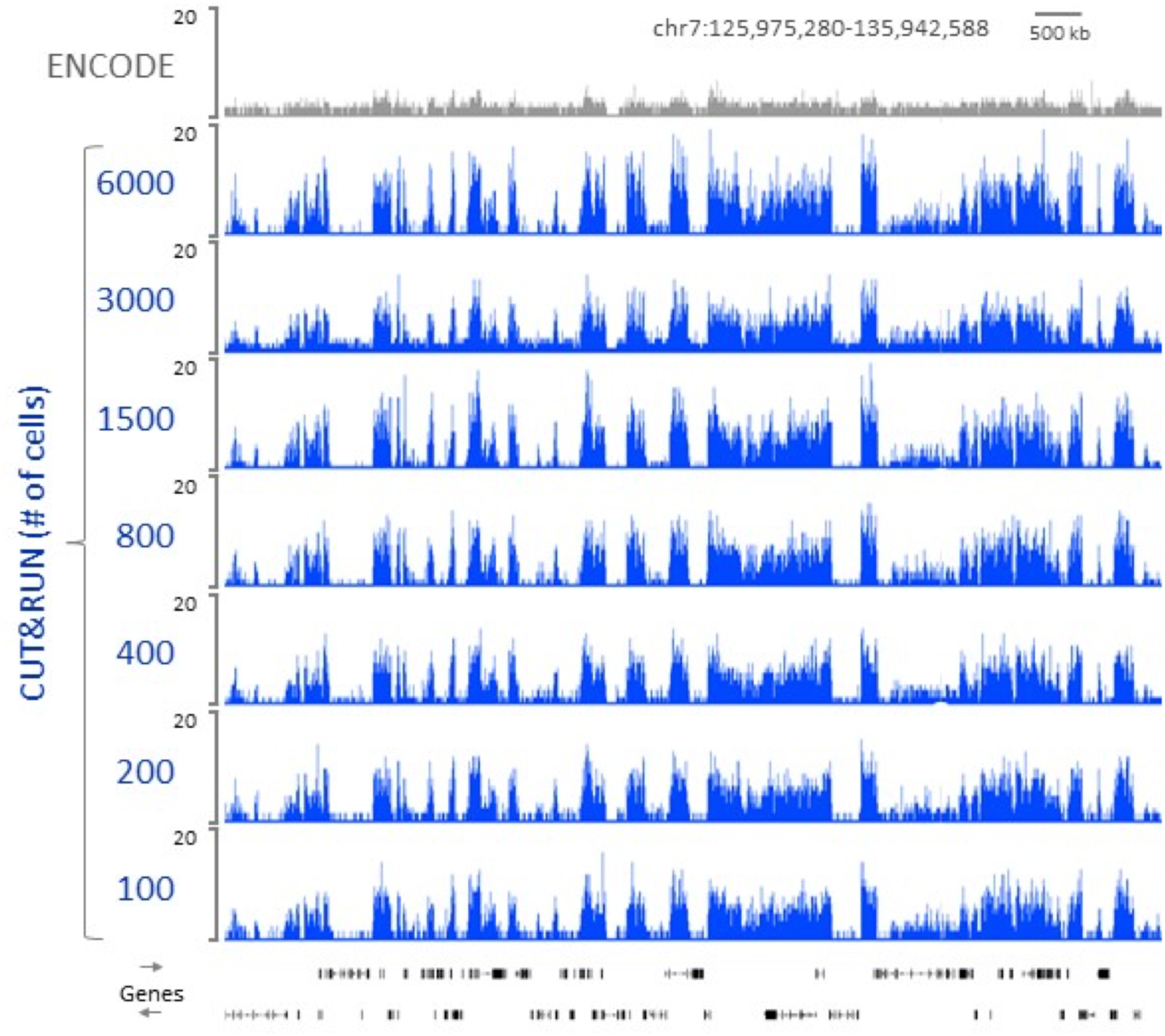
CUT&RUN of H3K27me3 requires only 100 cells to profile the human Polycomb chromatin landscape. A varying number of K562 cells was used as the starting material for profiling H3K27me3 by CUT&RUN. Following paired-end 25x25 bp Illumina sequencing and removal of duplicates, 7.5 million reads were randomly selected and used to generate bedgraphs representing raw counts, as indicated on the y-axis. For comparison, ENCODE X-ChIP-seq data (GSM733658) was similarly analyzed.

**Figure 4:**
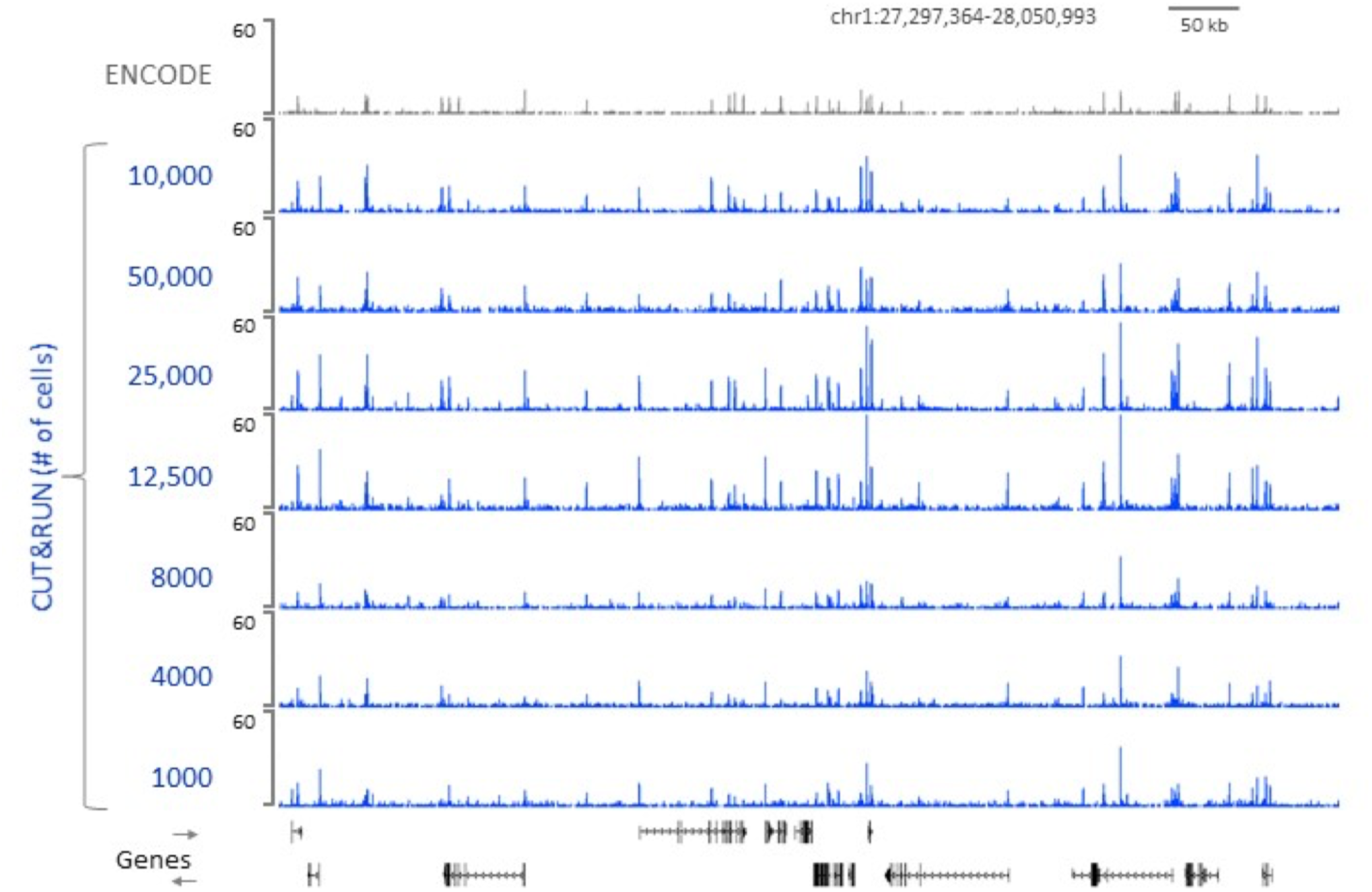
CUT&RUN requires only 1000 cells and 4 million reads to delineate human CTCF peaks. CUT&RUN was used to map CTCF binding sites in varying numbers of K562 cells. Following paired-end sequencing, 4 million non-duplicated reads were randomly selected and used to generate bedgraphs representing raw counts, as indicated on the y-axis. For comparison, ENCODE X-ChIP-seq data (GSM749690) was similarly analyzed.

Spin-column extraction (Steps 39-46) is simple and fast providing good recovery of fragments in the range of nucleosomes, while reducing the concentration of very large fragments that can interfere with library preparation (Supplementary Figure 4). Therefore, this DNA extraction option is preferred for most applications of CUT&RUN. However, for CUT&RUN of TFs at low cell numbers, organic extraction (Steps 47-58) is preferred for better recovery of small fragments.

**Supplementary Figure S4:**
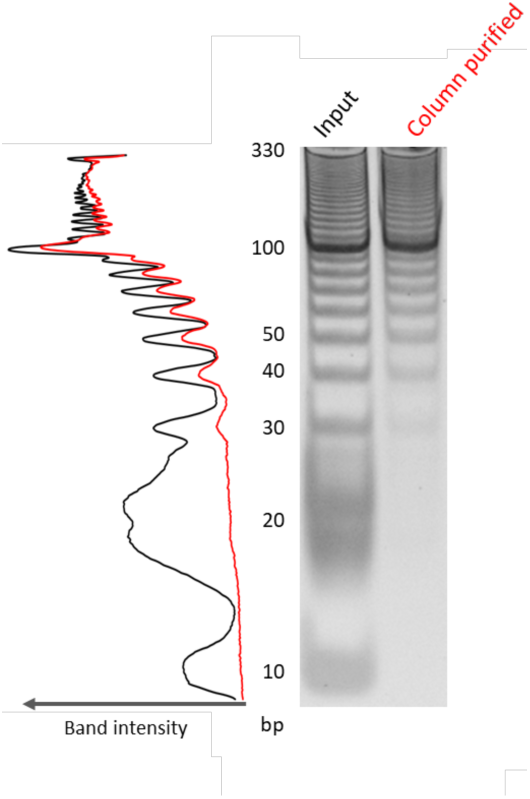
Spin-column DNA purification partially excludes both large and small fragments. To test the efficiency of spin columns in binding different length DNA fragments, 2 μg of 10 bp ladder was purified through the column and compared to 2 μg as input. DNA was resolved by 10% polyacrylamide gel electrophoresis and stained with SYBR-gold. Densitometry is shown on the left. For CUT&RUN, removal of large fragments reduces background, but removal of small fragments impacts recovery when profiling DNA-binding proteins. Therefore, spin-column purification (Steps 39-46) is preferred for nucleosomes, but might be less desirable for transcription factors and very low cell numbers, in which case the alternative PCI protocol (Steps 47-58) is recommended.

## AUTHOR CONTRIBUTIONS

P.S. and S.H developed the protocol, performed the experiments, analyzed the data and wrote the manuscript.

## ACKNOWLEDGEMENTS

We thank Paul Talbert for helpful comments on the manuscript, Christine Codomo for preparing Illumina sequencing libraries and Jorja Henikoff for bioinformatics.

## REFERENCES

1. Solomon, M.J. & Varshavsky, A. Formaldehyde-mediated DNA-protein crosslinking: a probe for in vivo chromatin structures. Proc Natl Acad Sci U S A 82, 6470–4 (1985).

2. Johnson, D.S., Mortazavi, A., Myers, R.M. & Wold, B. Genome-wide mapping of in vivo protein-DNA interactions. Science 316, 1497–502 (2007).

3. Barski, A. et al. High-resolution profiling of histone methylations in the human genome. Cell 129, 823–37 (2007).

4. Rhee, H.S. & Pugh, B.F. Comprehensive genome-wide protein-DNA interactions detected at single-nucleotide resolution. Cell 147, 1408–19 (2011).

5. Skene, P.J. & Henikoff, S. A simple method for generating high-resolution maps of genome-wide protein binding. eLife 4, e09225 (2015).

6. Teytelman, L., Thurtle, D.M., Rine, J. & van Oudenaarden, A. Highly expressed loci are vulnerable to misleading ChIP localization of multiple unrelated proteins. Proceedings of the National Academy of Sciences of the United States of America 110, 18602–7 (2013).

7. Park, D., Lee, Y., Bhupindersingh, G. & Iyer, V.R. Widespread misinterpretable ChIP-seq bias in yeast. PloS one 8, e83506 (2013).

8. Jain, D., Baldi, S., Zabel, A., Straub, T. & Becker, P.B. Active promoters give rise to false positive ‘Phantom Peaks’ in ChIP-seq experiments. Nucleic Acids Res 43, 6959–68 (2015).

9. Baranello, L., Kouzine, F., Sanford, S. & Levens, D. ChIP bias as a function of cross-linking time. Chromosome Res 24, 175–81 (2016).

10. Meyer, C.A. & Liu, X.S. Identifying and mitigating bias in next-generation sequencing methods for chromatin biology. Nat Rev Genet 15, 709–21 (2014).

11. Crawford, G.E. et al. Genome-wide mapping of DNase hypersensitive sites using massively parallel signature sequencing (MPSS). Genome Res 16, 123–31 (2006).

12. Giresi, P.G., Kim, J., McDaniell, R.M., Iyer, V.R. & Lieb, J.D. FAIRE (Formaldehyde-Assisted Isolation of Regulatory Elements) isolates active regulatory elements from human chromatin. Genome Res 17, 877–85 (2007).

13. Auerbach, R.K. et al. Mapping accessible chromatin regions using Sono-Seq. Proc Natl Acad Sci U S A 106, 14926–31 (2009).

14. Kent, N.A., Adams, S., Moorhouse, A. & Paszkiewicz, K. Chromatin particle spectrum analysis: a method for comparative chromatin structure analysis using paired-end mode next-generation DNA sequencing. Nucleic Acids Res 39, e26 (2011).

15. Henikoff, J.G., Belsky, J.A., Krassovsky, K., Macalpine, D.M. & Henikoff, S. Epigenome characterization at single base-pair resolution. Proc Natl Acad Sci U S A 108, 18318–23 (2011).

16. Buenrostro, J.D., Giresi, P.G., Zaba, L.C., Chang, H.Y. & Greenleaf, W.J. Transposition of native chromatin for fast and sensitive epigenomic profiling of open chromatin, DNA-binding proteins and nucleosome position. Nat Methods 10, 1213–8 (2013).

17. Bernt, K.M. et al. MLL-rearranged leukemia is dependent on aberrant H3K79 methylation by DOT1L. Cancer Cell 20, 66–78 (2011).

18. van Steensel, B., Delrow, J. & Henikoff, S. Chromatin profiling using targeted DNA adenine methyltransferase. Nat Genet 27, 304–8 (2001).

19. Schmid, M., Durussel, T. & Laemmli, U.K. ChIC and ChEC; genomic mapping of chromatin proteins. Mol Cell 16, 147–57 (2004).

20. Skene, P.J. & Henikoff, S. An efficient targeted nuclease strategy for high-resolution mapping of DNA binding sites. Elife 6(2017).

21. Hu, Z. et al. Nucleosome loss leads to global transcriptional up-regulation and genomic instability during yeast aging. Genes & development 28, 396–408 (2014).

22. Orlando, D.A. et al. Quantitative ChIP-Seq normalization reveals global modulation of the epigenome. Cell Rep 9, 1163–70 (2014).

23. Zentner, G.E., Kasinathan, S., Xin, B., Rohs, R. & Henikoff, S. ChEC-seq kinetics discriminate transcription factor binding sites by DNA sequence and shape in vivo. Nature Communications 6, 8733 (2015).

24. Corces, M.R. et al. Lineage-specific and single-cell chromatin accessibility charts human hematopoiesis and leukemia evolution. Nat Genet 48, 1193–203 (2016).

25. Liu, X. & Fagotto, F. A method to separate nuclear, cytosolic, and membrane-associated signaling molecules in cultured cells. Sci Signal 4, pl2 (2011).

26. Adam, S.A., Marr, R.S. & Gerace, L. Nuclear protein import in permeabilized mammalian cells requires soluble cytoplasmic factors. J Cell Biol 111, 807–16 (1990).

27. Brind’Amour, J. et al. An ultra-low-input native ChIP-seq protocol for genome-wide profiling of rare cell populations. Nat Commun 6, 6033 (2015).

28. Porreca, G.J. et al. Multiplex amplification of large sets of human exons. Nat Methods 4, 931–6 (2007).

29. Cusanovich, D.A. et al. Multiplex single cell profiling of chromatin accessibility by combinatorial cellular indexing. Science 348, 910–4 (2015).

30. Ramani, V. et al. Massively multiplex single-cell Hi-C. Nat Methods 14, 263–266 (2017).

31. Lieberman-Aiden, E. et al. Comprehensive mapping of long-range interactions reveals folding principles of the human genome. Science 326, 289–93 (2009).

32. Tang, Z. et al. CTCF-Mediated Human 3D Genome Architecture Reveals Chromatin Topology for Transcription. Cell 163, 1611–27 (2015).

33. Mumbach, M.R. et al. HiChIP: efficient and sensitive analysis of protein-directed genome architecture. Nat Methods 13, 919–922 (2016).

34. Chen, Y.B., A. “TSA-Seq”: a novel proximity mapping approach for studying three dimensional genome organization and function. (2016).

35. Beagrie, R.A. et al. Complex multi-enhancer contacts captured by genome architecture mapping. Nature 543, 519–524 (2017).

36. Wahba, L., Costantino, L., Tan, F.J., Zimmer, A. & Koshland, D. S1-DRIP-seq identifies high expression and polyA tracts as major contributors to R-loop formation. Genes Dev 30, 1327–38 (2016).

37. Sanders, M.M. Fractionation of nucleosomes by salt elution from micrococcal nuclease-digested nuclei. J Cell Biol 79, 97–109 (1978).

38. Davie, J.R. & Saunders, C.A. Chemical composition of nucleosomes among domains of calf thymus chromatin differing in micrococcal nuclease accessibility and solubility properties. J Biol Chem 256, 12574–80 (1981).

39. Henikoff, S., Henikoff, J.G., Sakai, A., Loeb, G.B. & Ahmad, K. Genome-wide profiling of salt fractions maps physical properties of chromatin. Genome Res 19, 460–9 (2009).

40. Kasinathan, S., Orsi, G.A., Zentner, G.E., Ahmad, K. & Henikoff, S. High-resolution mapping of transcription factor binding sites on native chromatin. Nature methods 11, 203–9 (2014).

41. Zentner, G.E. & Henikoff, S. High-resolution digital profiling of the epigenome. Nat Rev Genet 15, 814–27 (2014).

42. Fan, X., Lamarre-Vincent, N., Wang, Q. & Struhl, K. Extensive chromatin fragmentation improves enrichment of protein binding sites in chromatin immunoprecipitation experiments. Nucleic acids research 36, e125 (2008).

43. Teytelman, L. et al. Impact of chromatin structures on DNA processing for genomic analyses. PloS one 4, e6700 (2009).

44. Chung, H.R. et al. The effect of micrococcal nuclease digestion on nucleosome positioning data. PLoS One 5, e15754 (2010).

